# Transcriptomic Analysis of the Ocular Posterior Segment Completes a Cell Atlas of the Human Eye

**DOI:** 10.1101/2023.04.26.538447

**Authors:** Aboozar Monavarfeshani, Wenjun Yan, Christian Pappas, Kenechukwu A. Odenigbo, Zhigang He, Ayellet V. Segrè, Tavé van Zyl, Gregory S. Hageman, Joshua R. Sanes

## Abstract

Although the visual system extends through the brain, most vision loss originates from defects in the eye. Its central element is the neural retina, which senses light, processes visual signals, and transmits them to the rest of the brain through the optic nerve (ON). Surrounding the retina are numerous other structures, conventionally divided into anterior and posterior segments. Here we used high-throughput single nucleus RNA sequencing (snRNA-seq) to classify and characterize cells in the extraretinal components of the posterior segment: ON, optic nerve head (ONH), peripheral sclera, peripapillary sclera (PPS), choroid, and retinal pigment epithelium (RPE). Defects in each of these tissues are associated with blinding diseases – for example, glaucoma (ONH and PPS), optic neuritis (ON), retinitis pigmentosa (RPE), and age-related macular degeneration (RPE and choroid). From ∼151,000 single nuclei, we identified 37 transcriptomically distinct cell types, including multiple types of astrocytes, oligodendrocytes, fibroblasts, and vascular endothelial cells. Our analyses revealed a differential distribution of many cell types among distinct structures. Together with our previous analyses of the anterior segment and retina, the new data complete a “Version 1” cell atlas of the human eye. We used this atlas to map the expression of >180 genes associated with the risk of developing glaucoma, which is known to involve ocular tissues in both anterior and posterior segments as well as neural retina. Similar methods can be used to investigate numerous additional ocular diseases, many of which are currently untreatable.

## INTRODUCTION

Cells of the neural retina sense light (photoreceptors), process visual signals (interneurons), and transmit them to retinal output neurons (retinal ganglion cells, RGCs), which pass the signals to the brain through the optic nerve (ON). The retina does not, however, act alone: it is surrounded by a complex set of ocular tissues, which are conventionally divided into those of the anterior and posterior segments (Fig. 1A). The anterior segment includes the cornea, iris, and lens, which ensure that focused light reaches the photoreceptors; and the ciliary body, ciliary muscle, trabecular meshwork, and Schlemm’s canal, which produce and drain aqueous humor that nourishes avascular structures, clears away waste, and maintains intraocular pressure (IOP) (1). The posterior segment includes, in addition to the neural retina, the ON and its specialized proximal region, the optic nerve head (ONH); the sclera and a specialized region, peripapillary sclera (PPS), that encircles the ONH; retinal pigment epithelium (RPE), which supports the health and function of photoreceptors; and the choroid/choriocapillaris, which is the vascular supply that supports photoreceptors and RPE (Fig. 1A) (2, 3).

**Fig. 1.**
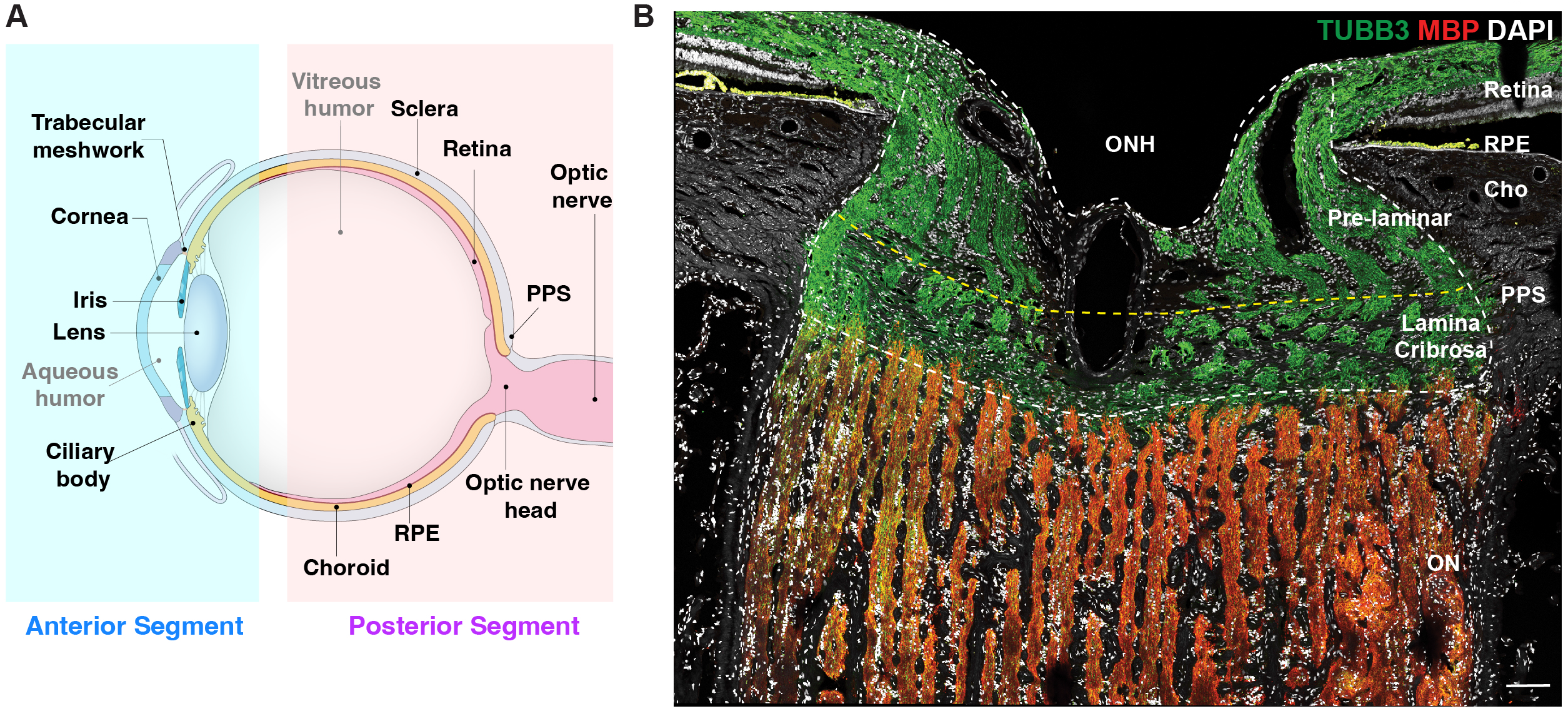
Anterior and posterior tissues of the human eye **(A**) Diagram of the human eye and the optic nerve, depicted in sagittal cross section. Structures represented in the ocular cell atlas are labeled. (**B**) Section of the optic nerve head (ONH) and surrounding tissues immunostained for myelin basic protein (MBP, red) and beta-tubulin (TUBB3, green). TUBB3 highlights bundles of axons in the retina and ONH and MBP highlights myelinating oligodendrocytes in the optic nerve (ON). Scale bar shows 100µm. PPS, Peripapillary Sclera; RPE, Retinal Pigment Epithelium; Cho, Choroid.

All of these structures have been implicated in conditions that impact vision. In the anterior segment, they include cataracts (lens), corneal dystrophies (cornea), uncorrected refractive error (cornea and lens), and anterior uveitis (iris/ciliary body), trachoma and onchocerciasis (cornea and conjunctiva) (4). Diseases of the posterior segment include optic neuritis, a frequent manifestation of multiple sclerosis (ON), posterior uveitis and scleritis (choroid/sclera), and diabetic retinopathy (retina) (5). Moreover, many blinding diseases involve multiple tissues – for example, trabecular meshwork, ONH, PPS and RGCs for glaucoma; photoreceptors, and RPE for retinitis pigmentosa; and photoreceptors, RPE and choroid for age-related macular degeneration (6–8). Thus, elucidating the pathogenic mechanisms underlying ocular diseases requires a comprehensive understanding of all ocular cell types and their interactions.

In previous studies, we used high-throughput single-cell and single-nucleus RNA sequencing (scRNA-seq and snRNA-seq) to catalog and molecularly characterize the cell types of the human anterior segment (9, 10). We and others also used scRNA-seq and snRNA-seq to characterize the adult human neural retina and RPE (11–19). Here, to complete a “v.1” cell atlas of the human eye, we used snRNA-seq to analyze the extra-retinal tissues of the posterior segment. We profiled ∼151,000 nuclei from the ON, ONH, PPS, RPE, choroid, and sclera, identified 37 cell clusters, and characterized the transcriptome of each. By dissecting tissues prior to profiling, we were able to better assess the tissue distribution of each cell type, identifying some types shared among tissues and others with striking tissue specificity.

Finally, to assess the expression of genes implicated in ocular disease, we focused on glaucoma, which is the leading cause of irreversible blindness worldwide (20). As noted above, multiple tissues are involved in this disease. The major modifiable risk factor for glaucoma is increased IOP, which is usually ascribed to defects in the outflow pathways of the anterior segment. Increased pressure leads to stress on the PPS and ONH, resulting in reactions of astrocytes and other cells in the lamina cribrosa (LC), a sieve-like structure in the ONH through which RGC axons pass (Fig. 1B) (21–23). These reactions, along with remodeling of the extracellular matrix, cause or exacerbate compression of RGC axons. The compression then leads to dysfunction and eventual death of the RGCs which is the proximate cause of glaucomatous vision loss (7, 21, 24). To date, at least 13 genes have been associated with early onset or congenital forms of glaucoma (21, 25–29). In addition, large GWAS meta-analyses have associated 127 genomic loci with primary open angle glaucoma (POAG; the most common late onset form of glaucoma), and 112 loci with IOP (30, 31). As an initial step in disentangling these interactions, we mapped the expression of >180 genes implicated in the pathogenesis of glaucoma or IOP regulation in the cells of the anterior and posterior segments.

## RESULTS

We profiled 36 tissue samples from 19 human donors with no history of ocular disease: 8 ON, 13 ONH, 4 PPS, 3 peripheral sclera, and 8 choroid/RPE (Fig. 2A, Table 1 and Table S1). Altogether, we recovered ∼210,000 single nucleus transcriptomes that we judged to be high quality based on the numbers of transcripts recovered and genes detected (Fig. S1). Initial analysis revealed that ∼60,000 of these transcriptomes were derived from neural retina present within all samples except those from ON; this was expected, given their proximity to other structures and limitations inherent to the manual dissection technique. We removed RPE (∼10,000) and retinal cells from the dataset. Neural retinal cells were not analyzed further and RPE was analyzed separately. We re-clustered the remaining ∼140,000 cells from all five tissue sources together (Fig. 2B).

**Fig. 2.**
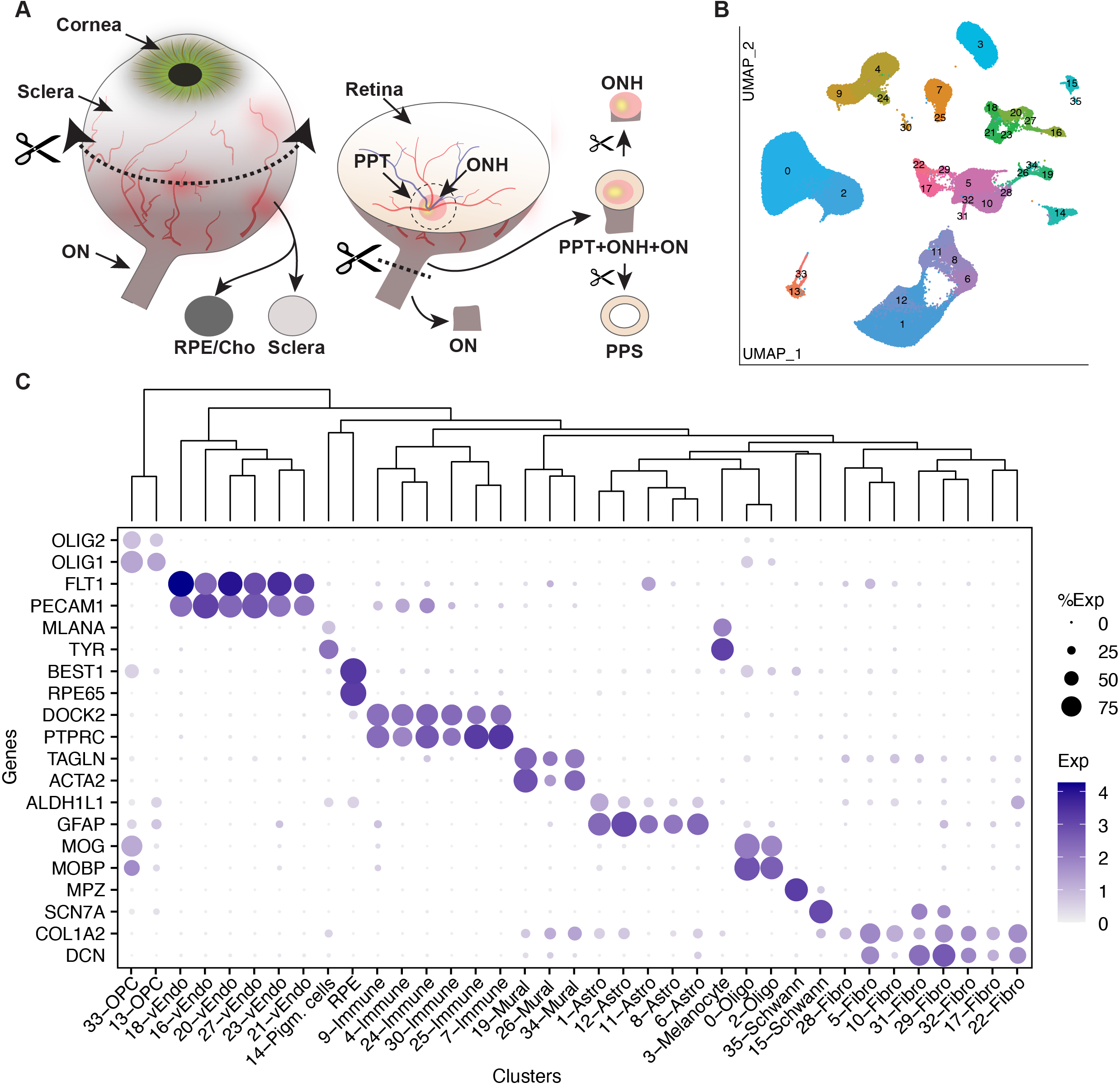
Collection of tissues from the posterior segment and initial transcriptomic analysis. (**A**) Schematic showing how tissues were dissected. Circumferential incision at the pars plana separated anterior and posterior segments, and tissues were dissected from the posterior segment using fine scissors and trephine tissue punches. (**B**) Clustering of single-nucleus expression profiles from all tissues visualized by Uniform Manifold Approximation and Projection (UMAP). Each tissue was processed separately. RPE and cells from neural retina were removed before the remaining ∼140k nuclei were pooled together to generate the UMAP. (**C**) Dot plot showing genes selectively expressed by each cell type. In this and subsequent figures, the size of each circle is proportional to the percentage of nuclei within a cluster expressing the gene and the color intensity depicts the average normalized transcript count in expressing cells. The dendrogram above the graph shows transcriptional relationships among cell types. ON, Optic Nerve; ONH, Optic Nerve Head; PPT, Peripapillary Tissues; PPS, Peripapillary Sclera; RPE, Retinal Pigment Epithelium; Cho, Choroid; Oligo, Oligodendrocytes; Endo, Vascular Endothelium; OPC, Oligodendrocyte Progenitor Cell.

**Table 1.** The number of cells and cell types that have been profiled and characterized in each ocular tissue of the human eye establishing the human eye atlas “v1”.

| **Tissue** | **#donors** | **#cells/nuclei** | **# types\*\*** |
| --- | --- | --- | --- |
| Cornea | 5 | 37485 | 7 |
| Lens | 5 | 13900 | 5 |
| Iris | 4 | 57422 | 9 |
| Ciliary body | 4 | 34132 | 10 |
| Trabecular meshwork | 4 | 39736 | 13 |
| Corneoscleral wedge | 4 | 36573 | 27 |
| Foveal retina | 5 | 56931 | 62 |
| Macular retina | 17 | 129775 | 98 |
| Peripheral retina | 10 | 55004 | 98 |
| Peripapillary retina | 17 | 59001 | 98 |
| Optic nerve head | 13 | 41178 | 32 |
| Peripapillary tissue | 4 | 4812 | 31 |
| Sclera | 3 | 4263 | 18 |
| Choroid | 8 | 36122 | 22 |
| RPE | 8 | 11422 | 1 |
| Optic nerve | 7 | 53603 | 16 |
| **Total** | **29** | **671359** |  |

\*\*types that composite $\geq 0.1\%$ of total cells from the tissue

Altogether, we identified 37 clusters, all derived from multiple samples (Table S2). All but RPE clusters are shown in a UMAP representation in Fig. 2B and numbered in the order of size (C0 has the most cells, C35 the fewest). We assigned them cell class labels based on selective expression of known markers (Fig. 2C) identifying 5 clusters as astrocytes, 2 each as oligodendrocytes, oligodendrocyte precursors, and Schwann cells, 8 as fibroblasts, 6 as vascular endothelium, 3 as mural (blood vessel-associated) cells, 6 as immune-related cells (microglia, macrophages and lymphocytes), and 3 as pigmented cells. As shown by a dendrogram representing transcriptomic similarity, types within a class were close transcriptomic relatives (Fig. 2C, top).

### Oligodendrocytes and their precursors

Two closely related clusters (C0 and C2) corresponded to mature oligodendrocytes. Both expressed markers of myelinating oligodendrocytes such as *MOBP* and *MOG* (Fig. 2C) (32, 33). They were, however, distinguishable by selective expression of multiple genes, including *ACTN2 and SLC4A8,* enriched in C0, and *LRRC7* and *DCC,* enriched in C2 (Fig. 3A). A comparison of the two oligodendrocyte populations to those described in human brain white matter by Jakël et al. (34) showed that C0 and C2 resembled their mature oligodendrocyte subtypes 1 and 5, respectively.

**Fig. 3.**
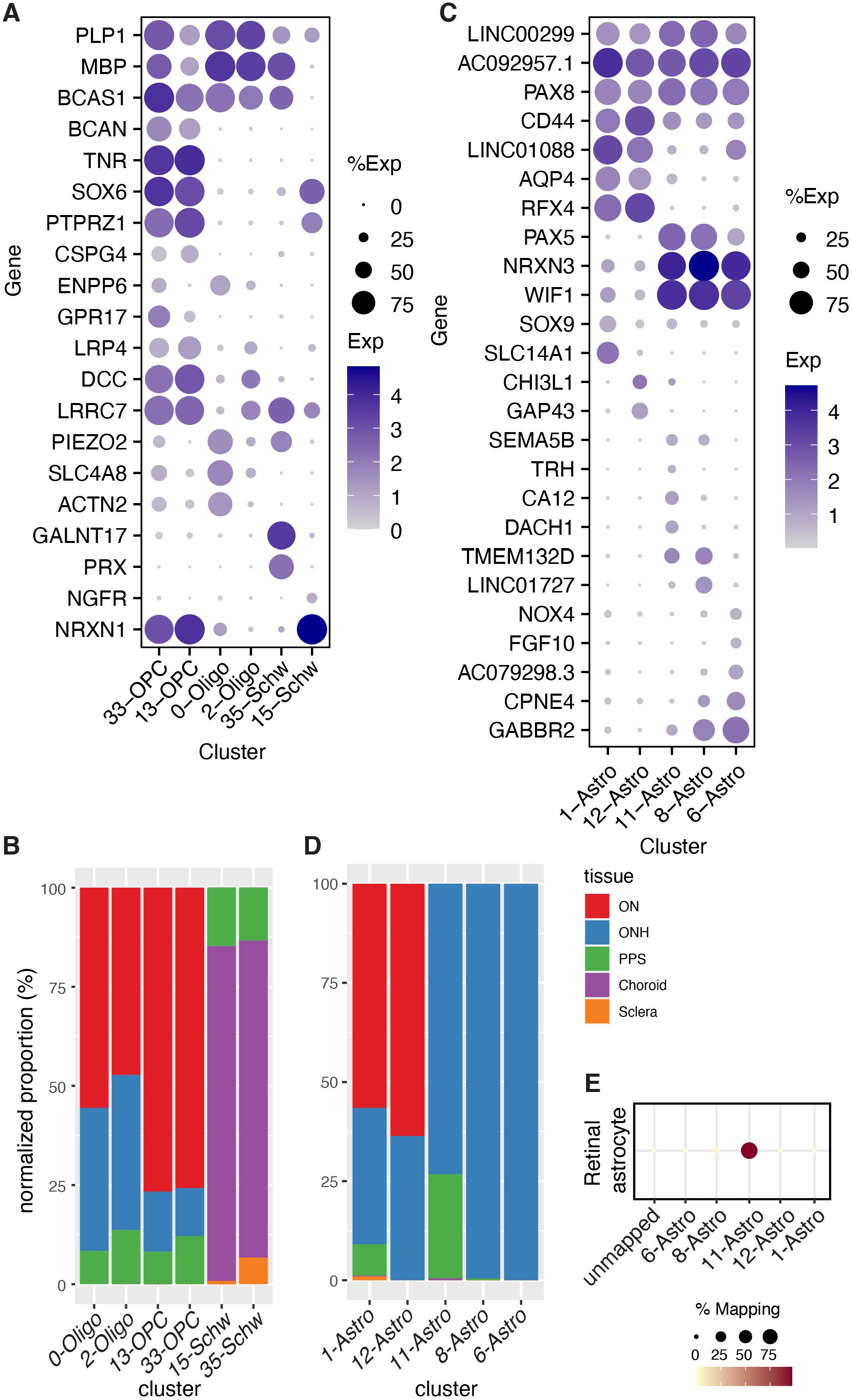
Glial cells. (**A**) Dot plot showing the expression of selected genes in oligodendrocyte, oligodendrocyte precursor cell (OPC), and Schwann cell clusters based on canonical markers (see Figure 2C). (**B**) Histogram showing the relative abundance of oligodendrocytes (Oligo), OPC, and Schwann clusters in five tissues. (**C**) Dot plot showing genes selectively expressed by all or some astrocyte types. (**D**) Histogram showing the relative abundance of each type of astrocyte in each tissue. (**E**) Transcriptional relationship of the astrocyte types identified in this study to retinal astrocytes profiled in van Zyl et al., (2022) (9). The color and size of each dot reflect the percentage of cells in retinal astrocytes (column) mapped to a corresponding type of astrocyte in ON/ONH (rows).

As expected, both types were most abundant in optic nerve samples and nearly absent from the sclera and choroid (Fig. 3B). However, 33% of C0 oligodendrocytes and 39% of C2 oligodendrocytes were derived from ONH samples, likely because those samples extended into the intraorbital portion of the ON (Fig. 2A). To test this idea, we stained sections of the optic nerve for myelin basic protein (MBP) which is an oligodendrocyte marker. Antibodies against this protein exclusively stained the myelinated portion of the ON (Fig. 1B), supporting the idea that oligodendrocytes recovered from ONH samples were derived from the ON. A small proportion of the C0 and C2 oligodendrocytes also appeared to be derived from the PPS samples, likely a result from the difficulty of completely separating the tightly apposed ONH and PPS during dissection.

Cells in the other two clusters (C13 and C33) expressed known markers of oligodendrocyte precursor cells (OPCs) such as *OLIG1* and *OLIG2* (33) (Fig. 2C). Although they were each other’s closest relatives, C33 was distinguishable from C13 by higher expression of several genes, including *GPR17, ENPP6*, and *PLP1*, which have been shown to be expressed by premyelinating oligodendrocytes (32) (Fig. 3A). Thus, C13 cells may be conventional OPCs while C33 may correspond to cells beginning to differentiate from OPCs to oligodendrocytes. Like oligodendrocytes, OPCs were most abundant in ON. OPCs were less abundant than mature oligodendrocytes in ONH (Fig. 3B), suggesting that they may be concentrated more distally along the ON.

### Schwann cells

Clusters C15 and C35 corresponded to Schwann cells, the glia of the peripheral nervous system. Both were predominantly derived from the choroid, with minor contributions from PPS and sclera (Fig. 3B). This distribution reflects the innervation of choroidal vessels by autonomic neurons (2). We identify C15 and C35 as comprising nonmyelinating and myelinating Schwann cells, respectively. C35 but not C15 expressed high levels of *PRX, MBP,* and *MPZ* which encode myelin proteins expressed by Schwann cells but not oligodendrocytes (Fig. 2C and 3A). In contrast, C15 is rich in *SCN7A, NCAM1, L1CAM*, and *NGFR* (Fig. 2C and 3A, Table S3), a profile known to represent nonmyelinating Schwann cells (35, 36).

### Astrocytes

Five clusters (C1, 6, 8, 11, 12) were close transcriptomic relatives and expressed known astrocyte markers including *GFAP,* and *ALDH1L1* (Fig. 2C). Additional markers that distinguished astrocytes from other cell classes included *PAX8,* and the non-coding RNAs *AC092957.1, LINC00299,* and *JAKMIP2-AS1* (Fig. 3C and S2A, and Table S3).

Comparisons amongst astrocyte clusters revealed distinct expression patterns and locations. The largest, C1, comprised ∼64% of all astrocytes. C1 and C12 were both most abundant in the optic nerve and expressed *AQP4* at high levels (Fig. 3C and D). They were present at far higher levels in the myelinated parts of the ON than ONH (Fig. S2B), consistent with a previous report on *AQP4* expression (37). Despite similarities in their transcriptomic profile and tissue distribution, they were molecularly distinct; C1 selectively expressed *SLC14A1* and *SOX9*, whereas C12 expressed higher levels of *GAP43* and *CHI3L1* (Fig. 3C).

Three clusters, C6, 8, and 11, were derived largely from ONH. All expressed *WIF1* and *PAX5* (Fig. 3C and S2D), but differentially expressed other genes that marked them as distinct types or states. For example, *DACH1* was expressed at the highest levels in C11, *LINC01727* in C8, and *FGF10* in C6 (Fig. 3C).

Whereas ∼99% of cells in C6 and C8 were derived from ONH, 26% of C11 cells were derived from PPS, after sample-size normalization (Fig. 3D). Based on transcriptomic similarity, we identified C11 as most similar to retinal astrocytes (12) (Fig. 3E) indicating that this cluster might be derived from retinal contamination, which was prominent in both ONH and PPS samples.

### Fibroblasts

Eight transcriptomically related clusters (C5, 10, 17, 22, 28, 29, 31, and 32) were classified as fibroblasts, based on their expression of known markers such as *COL1A1* and *DCN* (Fig. 2C) (38). This heterogeneity is consistent with numerous reports that have characterized multiple fibroblast types in single tissues (9, 39). The majority of cells in seven of the clusters were derived from a single tissue: C10 and 28 from choroid, C17, 22, 31 from PPS, C5 from sclera, and C29 from ONH; the eighth, C32, was derived to a similar extent from peripheral and peripapillary sclera (Fig. 4B and Table S4).

**Fig. 4.**
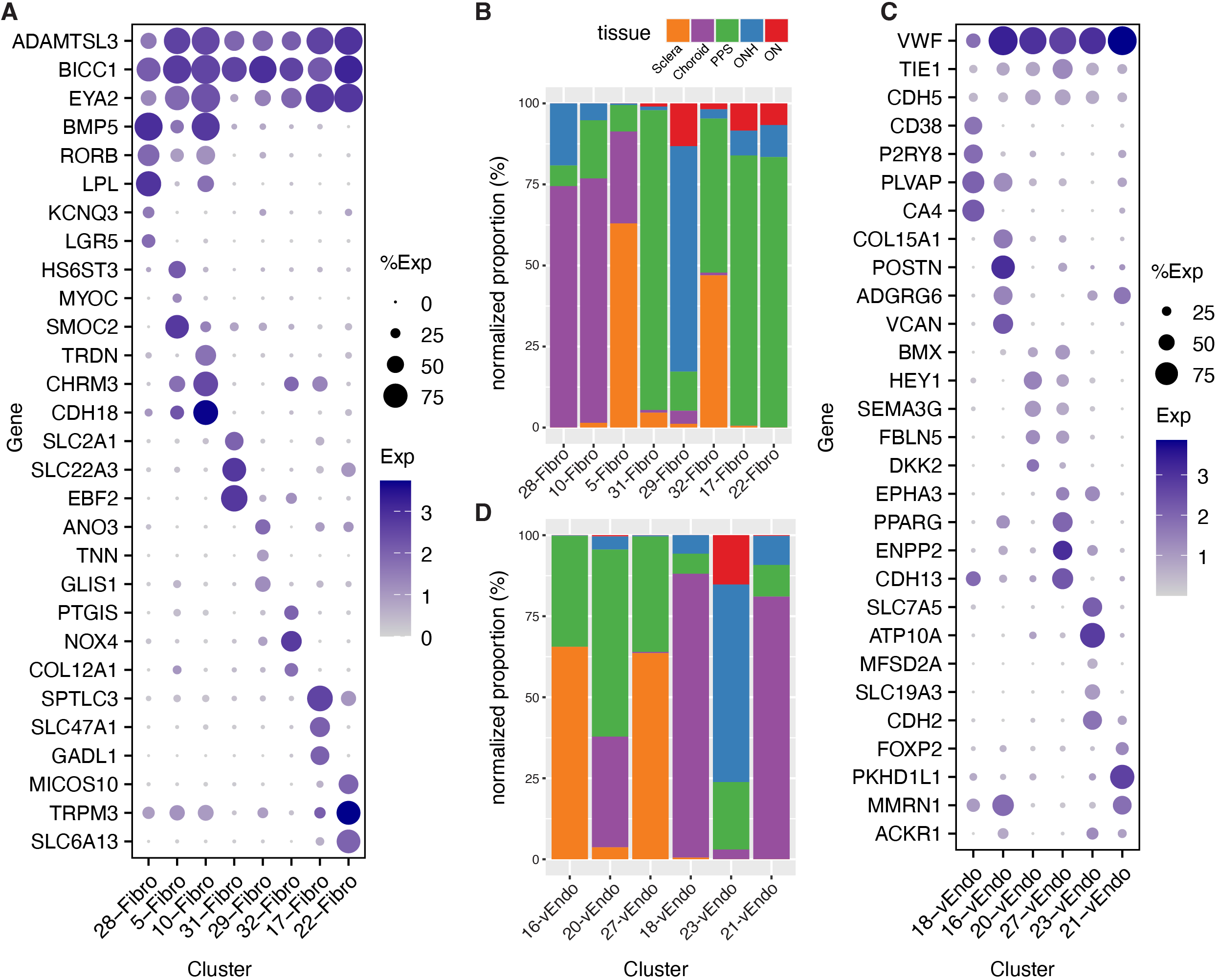
Fibroblasts and vascular endothelial cells. (**A**) Dot plot showing genes selectively expressed by all or each fibroblast cluster. (**B**) Histogram showing the relative abundance of each type of fibroblasts in each tissue. (**C**) Dot plot showing genes selectively expressed by all or each vascular endothelial cell type. (**D**) Histogram showing the abundance of each endothelial cell type in each tissue.

Differentially expressed genes included *HS6ST3* and *SMOC2* by C5; *TRDN* by C10; *SLC47A1* and *GADL1* by C17; *TRPM3*, and *SLC6A13* by C22; *KCNQ3* and *LGR5* by C28; *ANO3*, and *TNN* by C29; *SLC22A3*, and *EBF2* by C31; and *PTGIS* and *COL12A1* by C32 (Fig. 4A). Because fibroblasts are major sources of ECM, we mapped expression of ECM genes in these clusters, with emphasis on those reported to be present in the LC (Fig. S4B; see discussion). Comparison of posterior segment fibroblasts to those of the anterior segment (9) suggested that all except C31 were most similar to the anterior segment’s ciliary and/or scleral fibroblasts (Fig. S3A). This similarity may reflect the fact that the ciliary body and choroid are contiguous subdivisions of the uveal layer, and the scleral tissue encases the globe from its anterior-most aspect at the limbus to its posterior-most aspect encircling the optic nerve. C31 mapped to a type derived from the corneoscleral wedge, which we had termed “Fibro x” (9).

### Vascular endothelium

Six clusters (C16, 18, 20, 21, 23, and 27) were identified as vascular endothelial cells based on expression of markers such as *PECAM1* and *FLT1* (Fig. 2C). Using immunohistochemistry, we localized VWF+ endothelial cells in all tissues (Fig. S3B and C). C16 and C27 were derived predominantly from the sclera (including PPS); C18 and C21 were derived primarily from choroid; and C20 and C23 were derived from multiple tissues. C23 was the only cluster with a major contribution from ONH (>60%) and ON (>15%).

Based on expressed markers, we tentatively assign these cells to different vessel types. C16 cells expressed markers of venules including *ADGRG6, MMRN1*, and *POSTN*; C20 and C27 cells were enriched in the expression of artery/arteriole markers such as *SEMA3G, HEY1* and *BMX*; and C18 and C21 were enriched in the expression of genes previously shown to be associated with fenestrated capillaries, including *PLVAP* and *CA4* in C18, and post-capillary venules including *PKHD1L1* and *MMRN1* in C21 (38, 40–42). Mixed expression patterns in C21 and C23 (for example, expression of the venule marker, *ACKR1*) suggested that these cells might be derived from both venules and capillaries (42) (Fig. 4C and Fig S3C).

### Mural cells

Smooth muscle cells of the microvasculature, which primarily ensheath arterioles, and pericytes, which ensheath capillaries and venules, are collectively called mural cells (43). C19, 26, and 34 were identified as mural cells based on their expression of known markers such as *TAGLN* and *ACTA2* (40, 44–46) (Fig. 2C). Consistent with this assignment, *ACTA2*+ cells were closely associated with *PECAM*+ vascular endothelial cells (Fig. 5A and B).

**Fig. 5.**
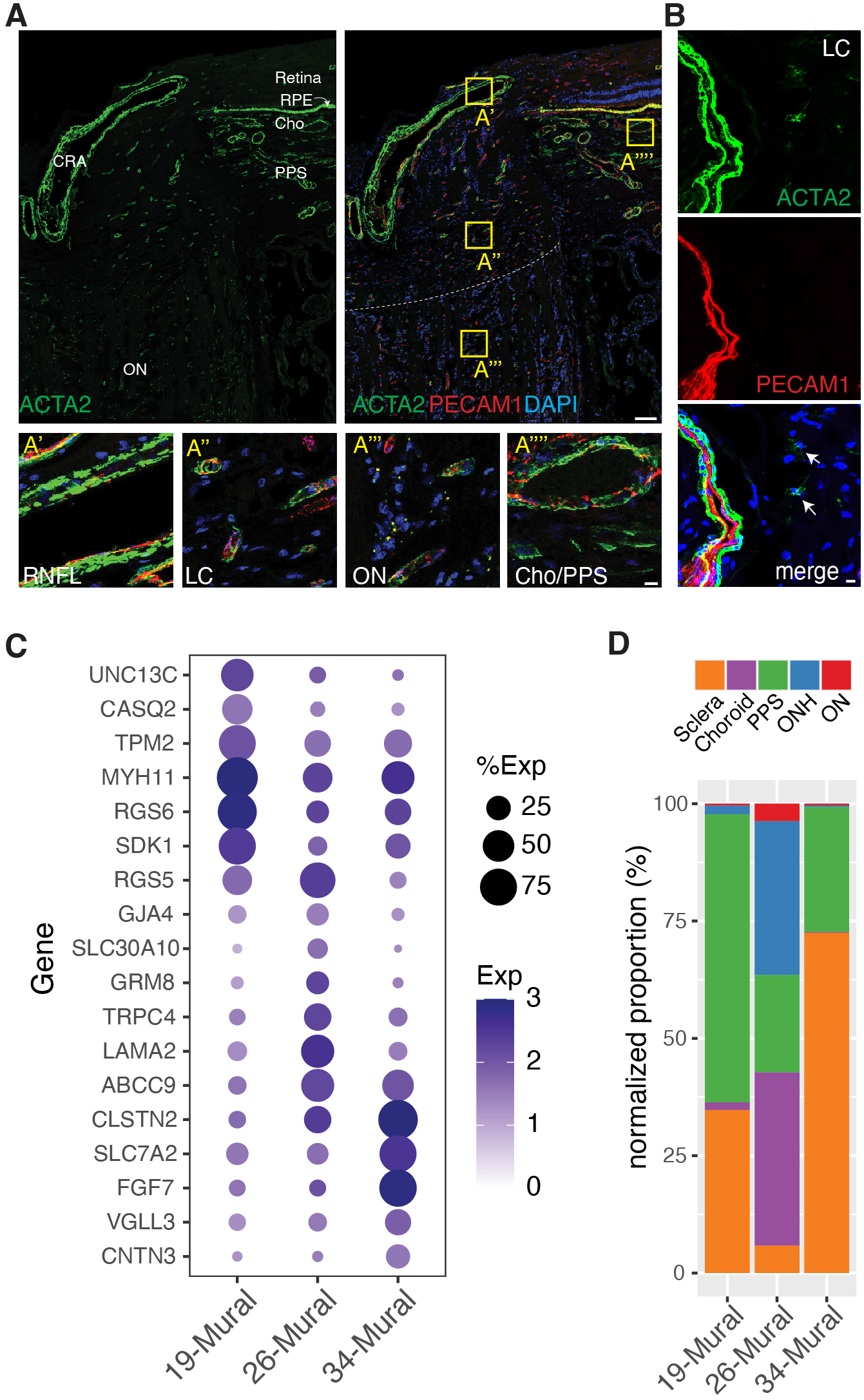
Mural cells. (**A and B**) Immunostaining for ACTA2 (green), and/or PECAM1 (red) in whole ONH (A) and LC region (B). Boxed areas in A are shown at higher magnification in lower panels. (**C**) Dot plot showing genes selectively expressed by cells in each mural cell cluster. (**D**) Histogram showing the abundance of the three mural types in each tissue. CRA, Central Retinal Artery; RNFL, Retinal Nerve Fiber Layer. Bars show 100µm in top panel A, 10µm in B and lower panel of A.

The three mural cell types were further distinguishable by selective expression of multiple genes including *UNC13C, RGS6*, and *CASQ2*, in C19, *TRPC4, RGS5*, and *LAMA2* in C26, and *FGF7, SLC7A2*, and *CLSTN2* in C34 (Fig. 5B). The expression profiles of the mural cells suggest that C19 and C34 cells are vascular smooth muscle cells, and C26 cells are pericytes. Consistent with these assignments, C19 and C34 were derived primarily from PPS and sclera, similar to that of the putative arteriolar endothelial cells, and C26 was derived in large part from choroid and ONH, similar to that of the putative capillary and venule-associated endothelial types (Fig. 4D and 5D). C26 corresponds to a *LAMA2+, TRPC4+* pericyte type we recently identified in the anterior segment (9) (Fig. 5C).

### Immune cells and microglia

Six clusters (C4, 7, 9, 24, 25, and 30) expressed markers of immune cells including *PTPRC* and *DOCK2* (47, 48) (Fig. 2C). Cluster 9 expressed numerous markers of microglia (e.g., *P2RY12*, and *ITGAM*; Fig. 6A). C4 and C24 expressed markers of macrophages (e.g., *MRC1*, and *CD163*) but could be differentiated from each other by selective expression of *IL1R2*, and *VCAN* in C24, and *CD163L1*, and *LYVE1* in C4 (49, 50) (Fig. 6A). We identified C7 as T lymphocytes (e.g., *CD2*+, and *THEMIS*+), C25 as NK cells (e.g., *KLRD1*+, and *KLRF1*+), and C30 as B lymphocytes (e.g., *EBF1*+, and *BCL11A*+) (51, 52) (Fig. 6A).

**Fig. 6.**
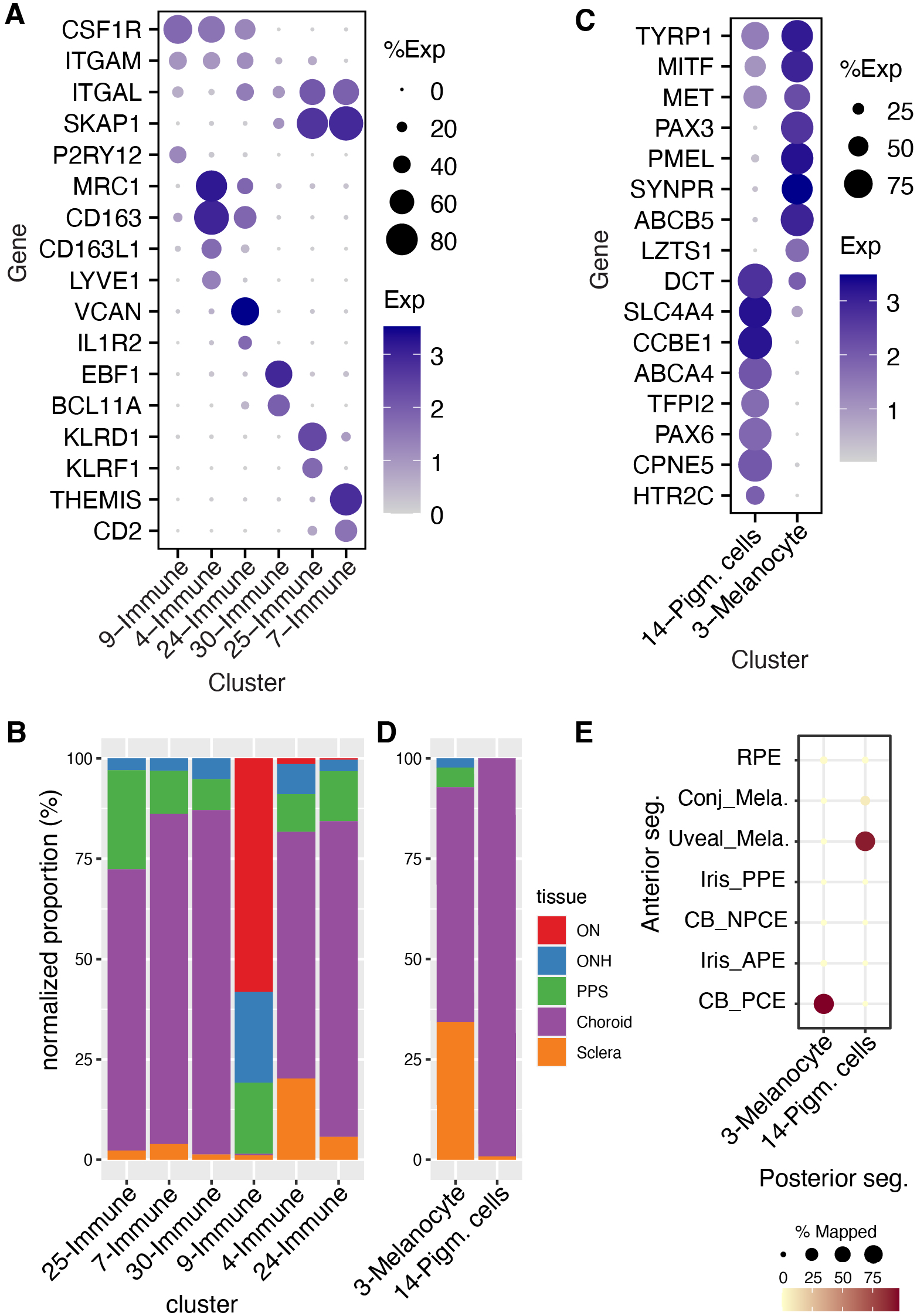
Immune and pigmented cells. (**A**) Dot plot showing genes selectively expressed by each of the immune-related cell types. (**B**) Histogram showing the relative abundance of each immune-related cluster in each tissue. (**C**) Dot plot showing genes selectively expressed by each type of melanocytes. (**D**) Histogram showing the abundance of the pigmented cell clusters C3 and C14 in each tissue. (**E**) Confusion matrix showing transcriptional correspondence of C3 and C14 in the current dataset to those in the anterior segment described in van Zyl et al., (9) and to RPE.

Most (80%) of the microglia were derived from ON and ONH, as expected from their known association with neural tissues. Consistent with this assignment, *AIF1*+ cells were abundant in retina, ONH and ON but rare in other tissues (Fig. S4C). In contrast, >95% of the other types were derived from peripheral and peripapillary sclera and choroid (Fig. 6B).

### Pigmented cells

Two cell types, C3, C14, expressed genes characteristic of pigmented cells such as *TYR* and *MLANA* (Fig. 2C). They could, however, be distinguished from each other by the expression of genes including *PMEL* and *PAX3* in C3 and *PAX6* and *CCBE1* in C14 (53) (Fig. 2C, 6C and S5). Almost all C14 cells and 71% of C3 cells were derived from choroid (Fig. 6D).

We identify C3 cells as melanocytes, based on their tissue source and expression pattern (Fig. 6, Table S3, and S4). Two types of melanocytes have been characterized in the anterior segment – a uveal type that resides in the iris and ciliary body, and a conjunctival type, that populates the conjunctiva (9). The C3 melanocytes were more closely related to the uveal than to the conjunctival type (Fig. 6E). C14 cells, in contrast, were more similar to pigmented epithelial cells of the ciliary body than to melanocytes (9), both in terms of overall transcriptomic similarity and differentially expressed genes (Fig. 6C, E). To our knowledge, cells of this type have not been described in the posterior segment. However, because they were obtained from peripheral samples, it is possible that they are derived from a transitional zone between the contiguous retinal pigment epithelium and ciliary body pigmented epithelial layer.

A third pigmented cell type, RPE, was derived largely from choroid/RPE samples, but was also present as contaminants in sclera and ONH samples. RPE pigment is generated by tyrosinase, but its levels are known to be low (54). The RPE dataset included >10,000 single nucleus transcriptomes from peripheral samples. Their transcriptomes generally resembled those reported in recent single cell studies (15, 17–19, 55–57), although our dataset is currently the largest. Along with known markers, we found several novel markers that are selectively expressed by RPE even when compared to all known ocular cell types (Fig. S5, Table S4).

We also profiled >1200 RPE nuclei from a macular sample and mapped genes differentially expressed between macula and periphery (Table S5). They include *WFDC1, CXCL14*, and *CALCB* enriched in macular RPE, and *PMEL* and *IGFBP5* enriched in peripheral RPE. Many of these genes were also reported to be differentially expressed in previous studies (15, 17–19, 55-58).

### Expression of glaucoma-associated genes

We previously generated cell atlases of the human neural retina and ocular anterior segment (9, 10, 12, 59). Together with the results presented above, they comprise a whole human eye cell atlas (Table 1).

A major motivation for generating this atlas was to map the expression of genes implicated in ocular diseases. As an illustration, we focus on glaucoma for two reasons. First, the most common form of glaucoma, POAG, is a leading cause of irreversible blindness worldwide (20). Second, as detailed in the introduction, cells and structures throughout the eye have been implicated in its pathogenesis. Over 200 genomic loci have been associated with POAG risk and/or IOP levels by GWAS analyses, together nominating >212 genes in the pathogenesis of glaucoma, increased IOP (which is easily measured) or both (25, 30, 31, 60). Here, we mapped the expression of the nearest gene/s to the GWAS loci for POAG risk or IOP regulation, along with rare Mendelian causes of POAG, in all ocular cell types that we have identified.

As shown in Fig. S6, at least some genes are expressed at significant levels in every ocular cell type, and most are expressed in multiple types. To seek expression patterns that correlate with disease features, we focused on cell types suspected to be involved in glaucoma, and assessed expression of four groups of genes: (1) Genes implicated in POAG susceptibility but not IOP regulation (“POAG only”). (2) Genes implicated in IOP regulation but not POAG susceptibility (“IOP only”). (3) Genes implicated in normal tension glaucoma, in which glaucoma progresses in the absence of increased IOP (“NTG”). (4) Genes associated with early onset or congenital glaucoma (“early onset”) (Table S6).

We calculated two disease gene scores: one that assesses the cell type specificity of each disease gene group in each cell type (z score, Fig. 7A), and another that evaluates the enrichment of genes from each disease group in each cell type compared to all glaucoma-related genes (ratio, Fig. 7B). For 14 of the 17 cell types (or small groups of related types), the two methods gave consistent results (Fig. 7). Genes in the early onset group, many being monogenic causes of large effect, were expressed at highest levels in the three main cell types of the outflow pathways, which we have called trabecular meshwork, iris and ciliary fibroblasts. (The names “ciliary” and “iris” reflect the presence of these types in adjacent structures as well as trabecular meshwork.)

**Fig. 7.**
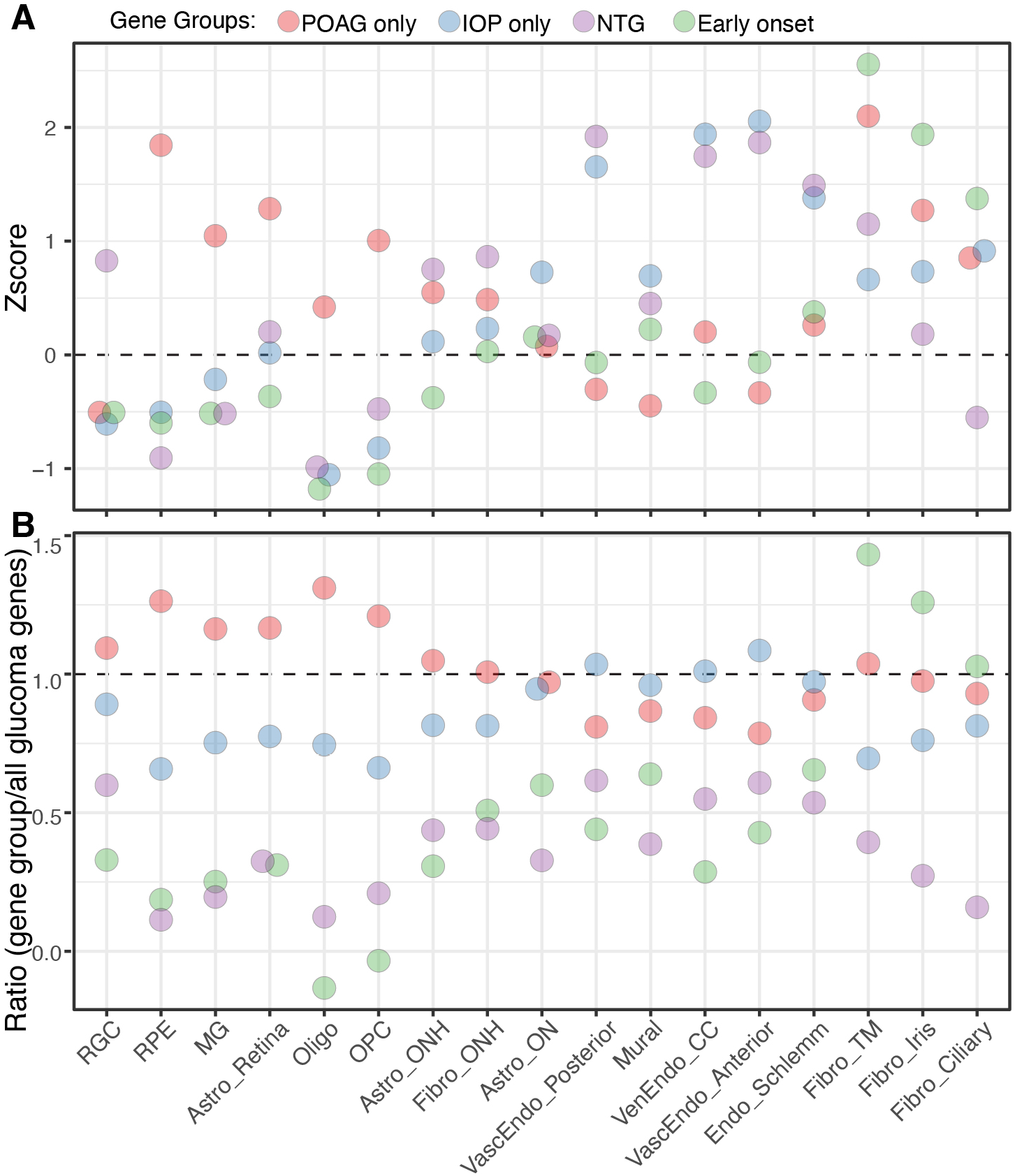
Expression of genes implicated in glaucoma. (**A**) Cell-type specific enrichment z-scores of groups of genes that have been associated with POAG but not IOP, IOP but not POAG, normal tension glaucoma (NTG), or early onset/congenital glaucoma. (**B**) The ratio of cell-type specific enrichment scores of each gene group to cell-type specific enrichment scores of all glaucoma associated genes (shown in Fig. S6B and table S6). CC, collector channel; Vasc, vascular; Ven, venous; Endo, endothelium.

Genes in the IOP only group demonstrated highest cell type specificity in vessel-associated cells (three vascular endothelial types and mural cells) as well as in the endothelium of Schlemm’s canal and in astrocytes of the optic nerve. Genes in the POAG only group were expressed at highest levels, compared to other gene groups, in retinal cells – retinal pigment epithelium, retinal astrocytes and Müller glia – as well as in oligodendrocytes, oligodendrocyte precursors and astrocytes of the optic nerve. Genes in the NTG group were also expressed at high levels in the endothelial types as assessed by z score. POAG only genes however demonstrated high cell type specificity in fibroblasts of trabecular meshwork.

For the three remaining cell types – RGCs, ONH astrocytes and ONH fibroblasts – the two methods give different results. When compared among cell types, the genes implicated in NTG exhibited cell type specificity in these three cell types. In contrast, when compared to expression of all glaucoma-related genes, genes in the POAG only group were expressed at the highest relative levels in all three cell types. The significance of these patterns is discussed below.

## DISCUSSION

Our study was motivated in large part by the knowledge that defects in the posterior segment have been implicated in the pathogenesis of many ocular diseases (see Introduction). However, although the posterior segment of primates, including humans, has been studied intensively using histological and histochemical methods (2, 61–66), knowledge of the cell types it contains and the genes that each type expresses remains incomplete. We therefore used a high-throughput RNA sequencing method to generate an atlas of cell types in six tissues of the posterior segment: ON, ONH, PPS, peripheral sclera, choroid, and RPE. In conjunction with our previous single cell analyses of anterior segment and neural retina (9, 10, 12, 59) and those of others (11-19, 55, 67-70), data presented here result in a first-pass cell atlas of the entire human eye (Table 1).

### Lamina Cribrosa

The LC region is a specialized region of the ONH that provides channels through which RGC axons enter the ON (21). Its two main compartments are bundles of axons that are unmyelinated proximal to the ON and instead ensheathed by astrocytes; and interstitial areas rich in extracellular matrix within which fibroblast-like cells (LC cells) are embedded (21, 23, 71). As a structure contiguous with the wall of the eye, it is exposed to different pressures from inside and outside the globe. Increased IOP leads to increased strain in the LC and the PPS with which it is continuous, leading to alterations in astrocytes, remodeling of the extracellular matrix, and possible deleterious effects on the vasculature that supplies the region (21). Normal tension glaucoma, in which there is no measurable abnormality of IOP, may result from increased susceptibility to the basal level of pressure (24).

Human LC cells have been tentatively identified in cultures prepared from ONH and studied intensively because of their importance in the pathogenesis of glaucoma (23, 64, 71–73). However, few markers have been found that label putative LC cells *in situ*, and we were unable to definitively annotate any cluster in our dataset as LC cells. Additionally, although previous reports proposed that ACTA2 is a marker for LC cells (71, 72), most ONH-derived ACTA2+ cells in the ONH are vessel-associated mural cells (pericytes and smooth muscle; Fig. 5A). Weakly ACTA2+ cells were also present (Fig. 5B), but they were predominantly present in the sclera (Fig. S4D).

We therefore assessed the distribution of cells in our dataset, based on the presumption that if LC cells comprised a unique population, we would expect them to be largely confined to the ONH or, given the likelihood of contamination, present at low abundance in the PPS. Three cell types stood out in this respect. Two were astrocytes (C6 and 8; Fig. 3D), which are, by definition, not LC cells. The third, fibroblast type C29, was enriched in, but not confined to ONH: nearly 70% of the cells in this cluster derived from ONH, with another ∼10% from PPS and ∼20% from ON (Fig. 4B).

Two additional results make C29 an intriguing candidate. First, following the suggestion of Paula *et al*. (23) that LC cells in the ONH share numerous structural and functional features with juxtacanalicular cells in the trabecular meshwork, we compared them to types characterized in our cell atlas of the anterior segment (9, 10). Indeed, C29 cells resembled ciliary fibroblasts, which are present in the trabecular meshwork as well as the ciliary body (Fig. S3A). Second, we compared the expression of genes in cell types of the ONH with those documented in cultured LC cells (71, 72, 74) (Fig. S4B). We relied on culture cells because of the paucity of analysis *in vivo*. C29 fibroblasts expressed most of the genes encoding known LC-associated molecules (Fig. S4B). Notably, however, in neither respect – gene expression or relationship to anterior segment – were C29 fibroblasts distinguished from those in other clusters.

Together, these results are consistent with two hypotheses. First, C29 fibroblasts may be the elusive LC cells. Alternatively, what have been referred to as LC cells may in fact be not a single type but rather a heterogeneous group that includes fibroblasts, pericytes, astrocytes and endothelial cells. Indeed, these cell types are likely to have been present in LC cultures and, tellingly, no studies of such cultures have shown that any gene or protein marks the entire population.

### Glaucoma

Genetic studies of POAG have identified numerous loci associated with disease risk, most of which are associated with increased IOP, which is a major risk factor for glaucoma (25, 30, 31, 60). We asked whether expression patterns of genes in these loci could provide insight into subsets of glaucoma, including normal tension and early onset (generally familial and monogenic) glaucoma. We also queried expression of two subsets of genes identified by GWAS: those associated with POAG but not increased IOP, and those associated with increased IOP but not POAG. Remarkably, genes in these four groups were expressed preferentially by different cell types.

Genes implicated in congenital glaucoma (e.g., *LTBP2, CYP1B1, PITX2*) were expressed at highest levels in three fibroblast types that are present in the iridocorneal angle and trabecular meshwork. This pattern is consistent with current understanding of the contribution of these cells to angle and outflow architecture.

Genes implicated in IOP but not POAG susceptibility were expressed at highest levels in vascular endothelial and mural cells as well as the endothelium of Schlemm’s canal. Together with the evidence that elevated IOP resulting from interference with outflow pathways leads to glaucoma, this pattern raises the question of whether some sources of increased IOP may be less damaging than others. Alternatively, these may be genes of small effect, so their signal in POAG GWAS fails to reach genome-wide significance.

Genes implicated in POAG susceptibility but not IOP regulation were expressed at highest relative levels in retinal cells – retinal pigment epithelium, retinal astrocytes and Müller glia – as well as in oligodendrocytes, oligodendrocyte precursors and astrocytes of the optic nerve. Although all three of the non-neural retinal cell types have been discussed as contributors to glaucoma pathogenesis (24, 75), mechanisms remain to be elucidated. Alterations in the ON in glaucoma have generally been viewed as consequences of axonal damage and loss, but our results support the hypothesis (76) that it may also play a causal role. The genes in this list provide entry points for further analysis.

Finally, genes implicated in POAG but not IOP were highly expressed in RGCs, ONH astrocytes and ONH fibroblasts by one metric (Fig. 7B), as were NTG by the other (Fig. 7A). These results are satisfying, in that dysfunction and death of RGCs are the cause of vision loss in glaucoma. Genes implicated in normal tension glaucoma were also enriched in endothelial cells, consistent with the idea that endothelial dysfunction can contribute to compression of RGC axons in the ONH (75).

In summary, our analysis of the cell types in which glaucoma susceptibility and causal genes are expressed supports long-held views of the importance of the trabecular meshwork and the ONH in glaucoma pathogenesis, but also draws attention to other cell types that have been less studied, such as astrocytes, oligodendrocytes and oligodendrocyte precursors in the ON. It will also be important to seek differences in gene expression between control and glaucomatous eyes. However, such changes could be either causes or results of disease processes. Therefore, knowing expression patterns in normal tissue of genes implicated in the disease is an essential complement to such studies.

## MATERIALS & METHODS

### Tissue Acquisition, Dissection, and Processing

Human eyes were obtained post-mortem at a median of 6 hours from death either from the Massachusetts General Hospital (MGH) via the Rapid Autopsy Program or through the Steele Center for Translational Medicine at the John A. Moran Eye Center (SCTM), University of Utah (Table S1). Whole globes were transported either to Harvard University or the University of Utah in a humid chamber on ice and processed within an hour of enucleation.

Dissection was performed under a microscope. To isolate the anterior and posterior segments, a surgical blade was used to make a small stab incision ∼4 mm posterior to the limbus at the pars plana. The incision was extended circumferentially with curved scissors to yield a separated anterior segment and posterior segment. Each segment was placed in a petri dish filled with Ames’ medium equilibrated with 95% O2/5% CO2 or sterile PBS, on ice.

A 4mm trephine punch centered around the optic disc was obtained from the posterior segment. It contained ONH, peripapillary sclera (PPS), choroid, retinal pigment epithelium (RPE) and retina, as well as the proximal ON. In a few cases, peripapillary tissue was removed and the ON stump were cut at the level of sclera to exclude intraorbital ON portion. ON beyond the ONH was dissected from surrounding dura and arachnoid mater and cut into 4mm long pieces to yield ON. Dissected tissues were immediately placed along the wall of a cryogenic vial, ensuring minimal liquid was present, and submerged in dry ice or liquid nitrogen. For long-term storage, the cryogenic vials were moved to a -80°C freezer. Samples collected at the University of Utah were shipped to Harvard University on dry ice.

For single-nuclei isolation, frozen tissue samples were homogenized in a Dounce homogenizer in 1ml lysis buffer (Tris, CaCl2, MgCl2, NaCl, RNase inhibitor, DNase I, and 0.1% NP-40) and passed through a 40-µm cell strainer. The filtered nuclei were then pelleted at 500 rcf for 5 min and resuspended in 2% BSA with DAPI counterstain for cell sorting on a flow cytometer. The sorted nuclei were pelleted again at 500 rcf for 5 min, resuspended in 0.04% non-acetylated BSA/PBS solution, and adjusted to a concentration of 1000 nuclei/µL. The integrity of the nuclear membrane and presence of non-nuclear material were assessed under a brightfield microscope before loading into a 10X Chromium Single Cell Chip (10X Genomics, Pleasanton, CA) with a targeted recovery of 8000 nuclei.

Single nuclei libraries were generated using the Chromium 3’ V3, or V3.1 platform (10X Genomics, Pleasanton, CA) following the manufacturer’s protocol. Briefly, single nuclei were partitioned into Gel-beads-in-EMulsion (GEMs) where nuclear lysis and barcoded reverse transcription of RNA would take place to yield full-length cDNA; this was followed by amplification, enzymatic fragmentation and 5’ adaptor and sample index attachment to yield the final libraries.

### RNA Sequencing and Data Analysis

Libraries were sequenced on an Illumina NovaSeq at the Bauer Core Facility at Harvard University. Sequencing data were demultiplexed and aligned using Cell Ranger software (version 4.0.0, 10X Genomics, Pleasanton, CA). The human genome reference file GRCh38 and its associated transcriptome file (release 101) were downloaded from Ensembl. The transcriptome file was modified before the alignment so that each “transcript” entry was converted to the “exon” entry to account for pre-mRNA. Downstream analysis followed the same pipeline described previously (9, 12), utilizing the R package “Seurat V4” (77). In summary, nuclei collected from ONH, ON, PPS and sclera were analyzed separately without batch correction. After clustering, clusters formed by retinal cells including photoreceptors, horizontal cells, bipolar cells, amacrine cells, retinal ganglion cells, and Müller glia were identified based on the expression of canonical markers and removed from further analysis. RPE cells were also removed and analyzed separately. The remaining cells were pooled together, and re-clustered. At each step, the count matrix was normalized and scaled. Batch correction was performed using the “IntegrateData” function. Unsupervised clustering was performed using the “FindClusters” function with “Louvain algorithm with multilevel refinement” as the modularity optimization algorithm. For comparison of types from current study to those identified in the anterior segment, the extreme gradient boosting algorithm utilizing the R package “XGBoost” were applied to the training dataset (types from the anterior segment), and the generated classifier was then used to make predictions on individual cells in the current dataset. To display the tissue distribution of each cell type (histograms in Fig. 3-6), we calculated the fraction of cells obtained from each tissue, and then corrected the values to account for sample size differences (Table S4).

### Glaucoma gene expression analysis

After cell type annotation, the expression of four gene sets (Table S6) relevant to glaucoma (30, 31) were assessed as described in van Zyl et al (9). The sets are defined in the Results. For mapping the expressions of ocular disease related genes, the POAG gene list was taken from Supplementary Data 4 in Gharahkhani et al (30), and the IOP list from Khawaja et al (31), which contain the nearest gene/s to 127 POAG and 133 IOP lead GWAS variants, respectively. Only genes that were expressed in at least 5% of cells in one or more clusters were included. The expression scores were calculated by first subtracting the averaged expression of all genes from the averaged expression of genes of individual sets in each cell type. To assess the cell types that selectively expressed genes associated with each disease group, we subtracted the mean value across all cell types from the raw score for each cell type, and divided by the standard deviation of values for all cell types, thus generating a z-score (Fig. 7A and Fig. S6). To evaluate the enrichment of genes from each disease group in each cell type, we divided the raw score of each group by that for all glaucoma-related genes in the cell type (Fig. 7B and Fig. S6). For the bubble plot, the gene scores were normalized across cell types for each set.

### Immunohistochemistry and Imaging

Whole globes were fixed in 4% paraformaldehyde (in PBS) for 2-24 hr and then transferred to PBS. ONH tissues (4mm punches) were dissected out and placed in 30% sucrose in PBS overnight at 4°C, before being embedded in tissue freezing medium and mounted onto coated slides in 20 μm coronal sections. Slides were incubated for 1 hour with 5% donkey serum (with 0.1% TritonX) at room temperature, then for overnight with primary antibodies at 4°C, and finally for 2 hours with secondary antibodies at room temperature. Antibodies used are listed in Table S7. Images were acquired on Zeiss LSM 710 confocal microscopes with 405, 488-515, 568, and 647 nm lasers, processed using Zeiss ZEN software suites, and analyzed using ImageJ (NIH). Images were acquired with 20X lenses at the resolution of 1024×1024 pixels, a step size of 0.5-1.5μm, and 90μm pinhole size.

## ACKNOWLEDGEMENTS

This work was supported by the Chan-Zuckerberg Initiative (CZF-2019-002459), NIH (5K12EY016335, EY028633 and U01 MH105960), an unrestricted grant from Research to Prevent Blindness to the Department of Ophthalmology and Visual Sciences, University of Utah, and by charitable donations to the Sharon Eccles Steele Center for Translational Medicine. A.M. was supported by T32NS007473, F32EY031930, and K99EY033457. We thank Lisa Nichols and Jennifer Mohler for assistance in acquiring and processing ocular tissues, and the tissue donors and their families for their generosity.

## DATA AVAILABILITY

The accession number at Gene Expression Omnibus for the raw data reported in this paper is GSExxxxxx. (Deposition is in progress. Accession number will be supplied when available. In addition, all data will be uploaded to the Broad Single Cell Portal.

## TABLE AND SUPPLEMENTARY TABLE LEGENDS

Supplementary Table 1. Information about human donor eye tissue processed for snRNA-seq. Hu, human; OD, right eye; OS, left eye; MGH, Massachusetts General Hospital; COD, cause of death; DTP, death to processing time.

Supplementary Table 2. The number of high-quality single nucleus transcriptional profiles obtained for each donor, tissue and cluster.

Supplementary Table 3. Differentially expressed genes in each cluster compared with all other clusters.

Supplementary Table 4. The number, proportion and normalized proportion of single nucleus transcriptional profiles obtained for each cluster.

Supplementary Table 5. Differentially expressed genes between macular and peripheral RPEs. Highlighted genes (in blue) indicate differentially expressed genes that have been reported in previous literature.

Supplementary Table 6. A list of all glaucoma associated genes (grouped as “IOP only”, “POAG only”, “NTG”, “Early Onset”, and “POAG and IOP”) used to calculate scores in in Figs. 7 and S6A.

Supplementary Table 7. Antibodies used for histological studies.

**Fig. S1.**
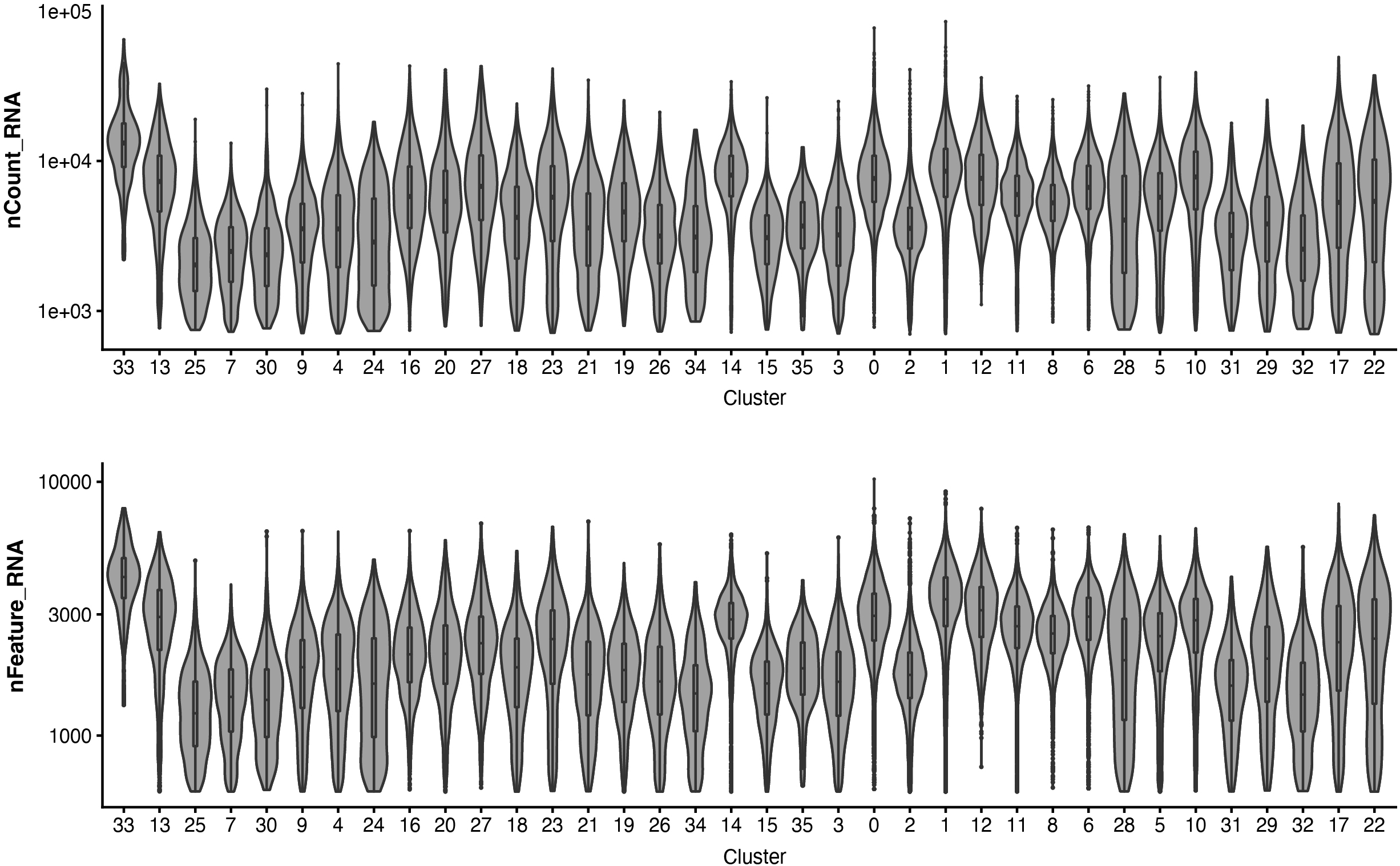
Violin plot showing the number of transcripts and genes detected in each cluster.

**Fig. S2.**
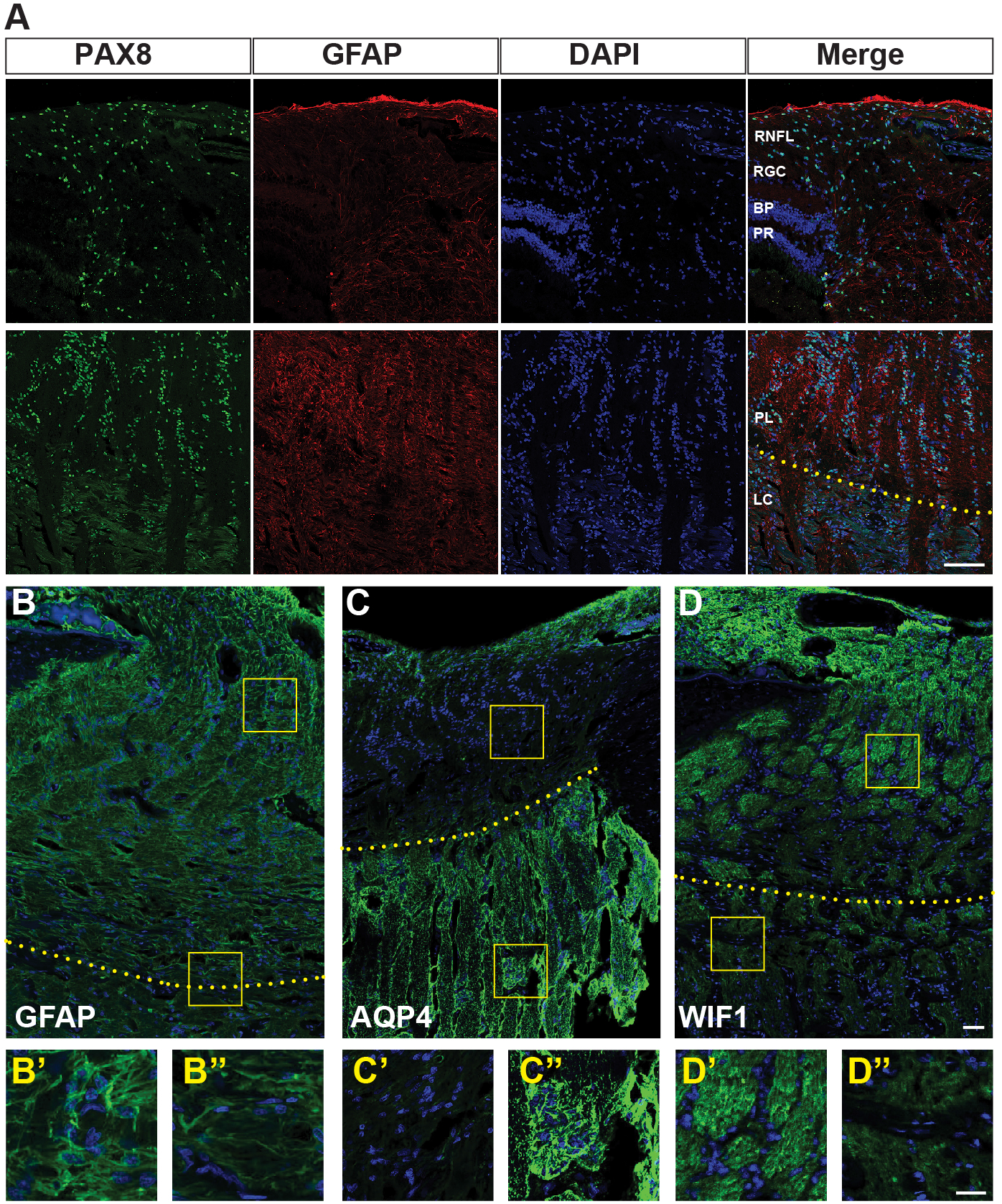
(**A**) Immunostaining for PAX8 (as a novel pan marker of astrocytes) and GFAP showing distribution of PAX8+ astrocytes in two different regions of ONH tissue. (**B-D**) Immunostaining for GFAP(B), AQP4 (C), and WIF1 (D) highlights heterogeneity of astrocytes in ONH and ON tissues. Boxed areas in top panels are shown at higher magnification in lower panels. Yellow dotted line indicates the upper (in A) and lower (in B-D) border of LC. Bars show 100µm in B-D, and 50µm in magnified boxes B’-D’’. RGC, Retinal Ganglion Cells; BP, Bipolars, PR, Photoreceptors; RNFL, Retinal Nerve Fiber Layer; LC, Lamina Cribrosa; PL, Prelaminar region.

**Fig. S3.**
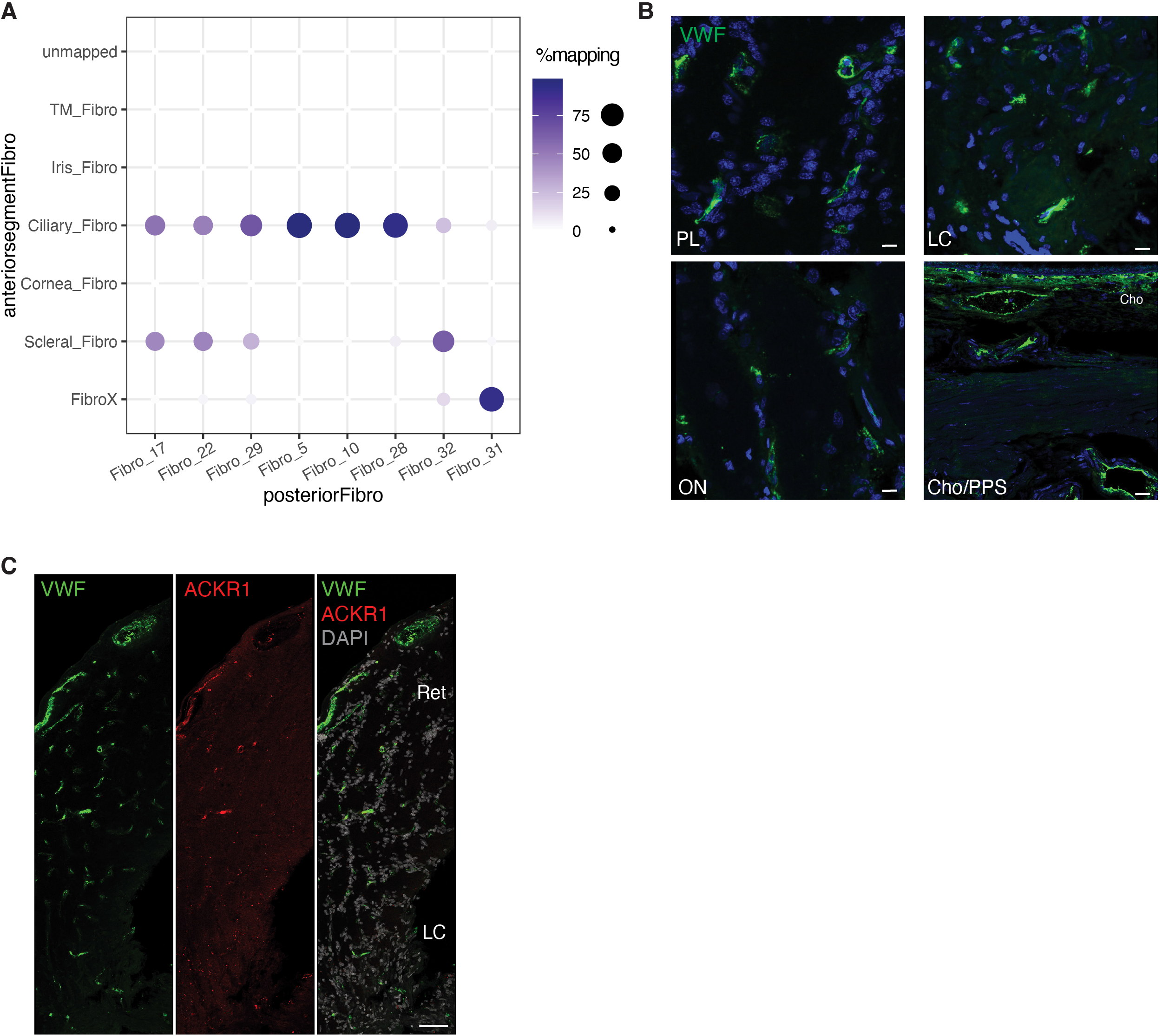
(**A**) Confusion matrix showing transcriptional correspondence of fibroblasts in the current dataset to those in the anterior segment (van Zyl et al., 2022). (**B**) Immunostaining for VWF in prelaminar region of the ONH (PL), lamina cribosa (LC), optic nerve (ON), and peripapillary sclera (PPS). (**C)** Immunostaining for VWF (green) and ACKR1 (red) in human ONH. Bars show 10µm in B and 100µm in C.

**Fig. S4.**
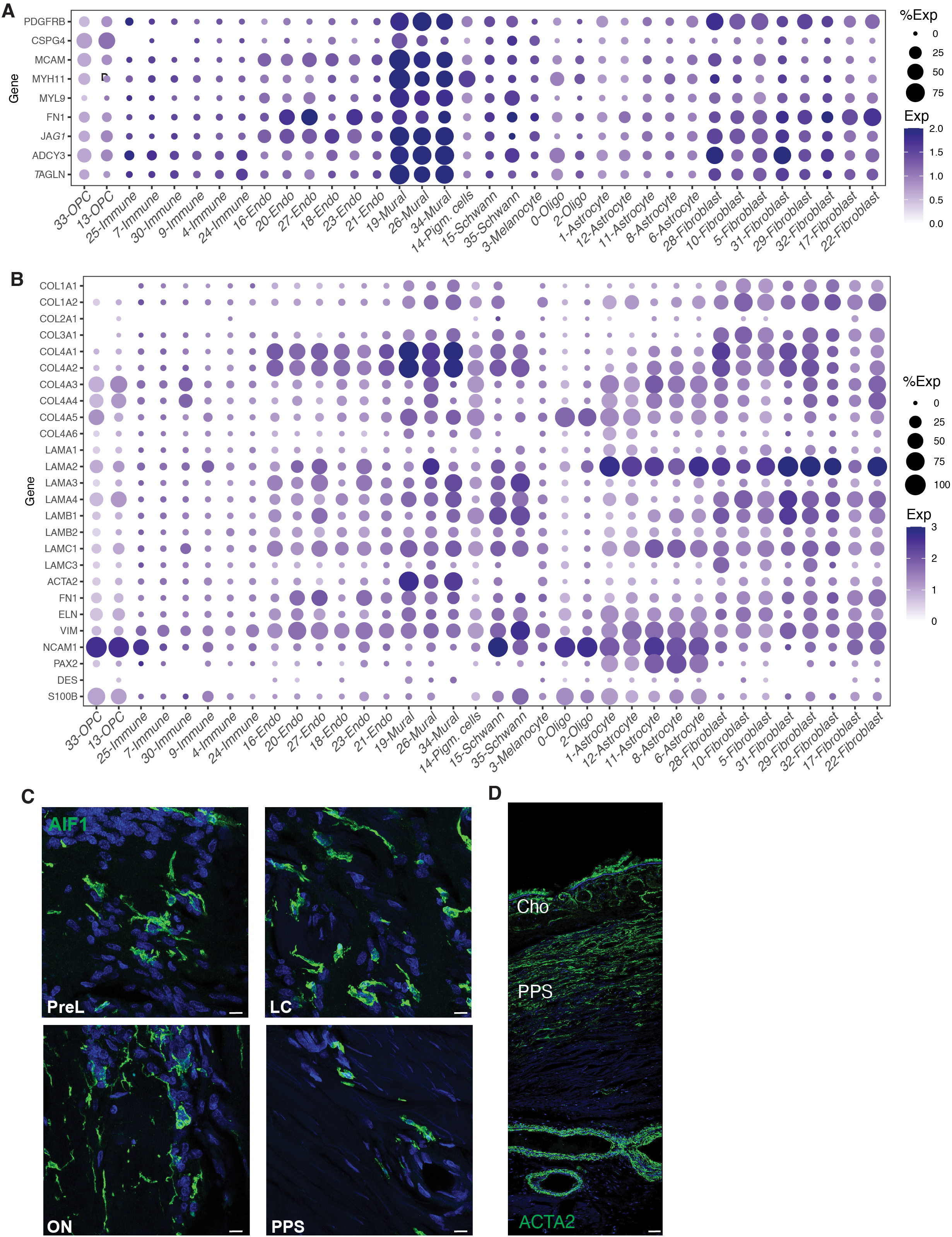
(**A**) Dot plot showing expression of reported mural cell markers. (**B**) Dot plot showing expression of ECM genes and genes reported to be expressed by LC cells in all clusters. (**C**) Immunostaining for AIF1/IBA1 showing microglia and/or macrophages in ONH, ON, and scleral tissues. (**D**) Immunostaining for ACTA2 (green) in PPS. Bars show 10µm in C and 50µm in D.

**Fig. S5.**
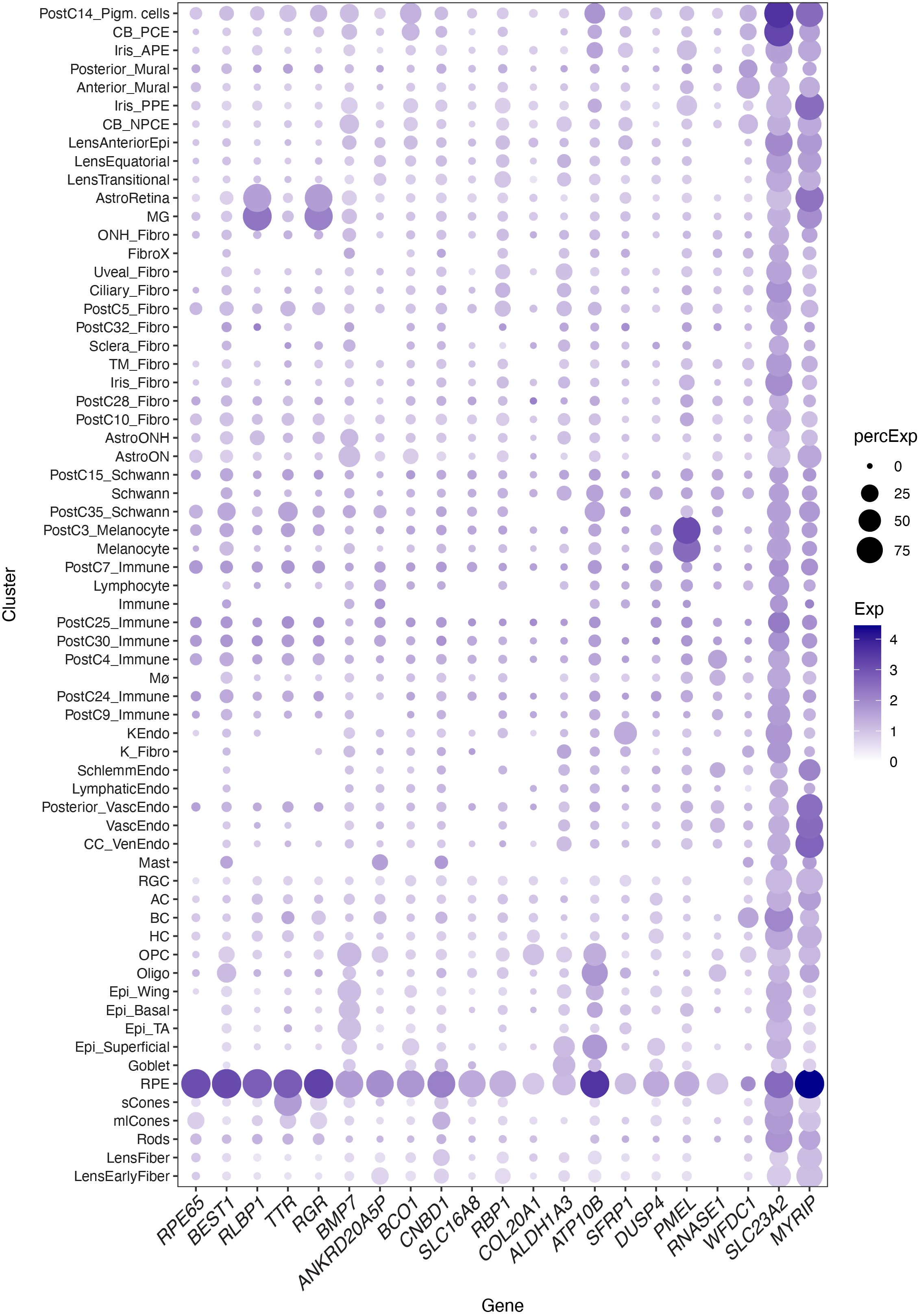
Dot plot showing expression of known and novel marker genes for RPE.

**Fig. S6.**
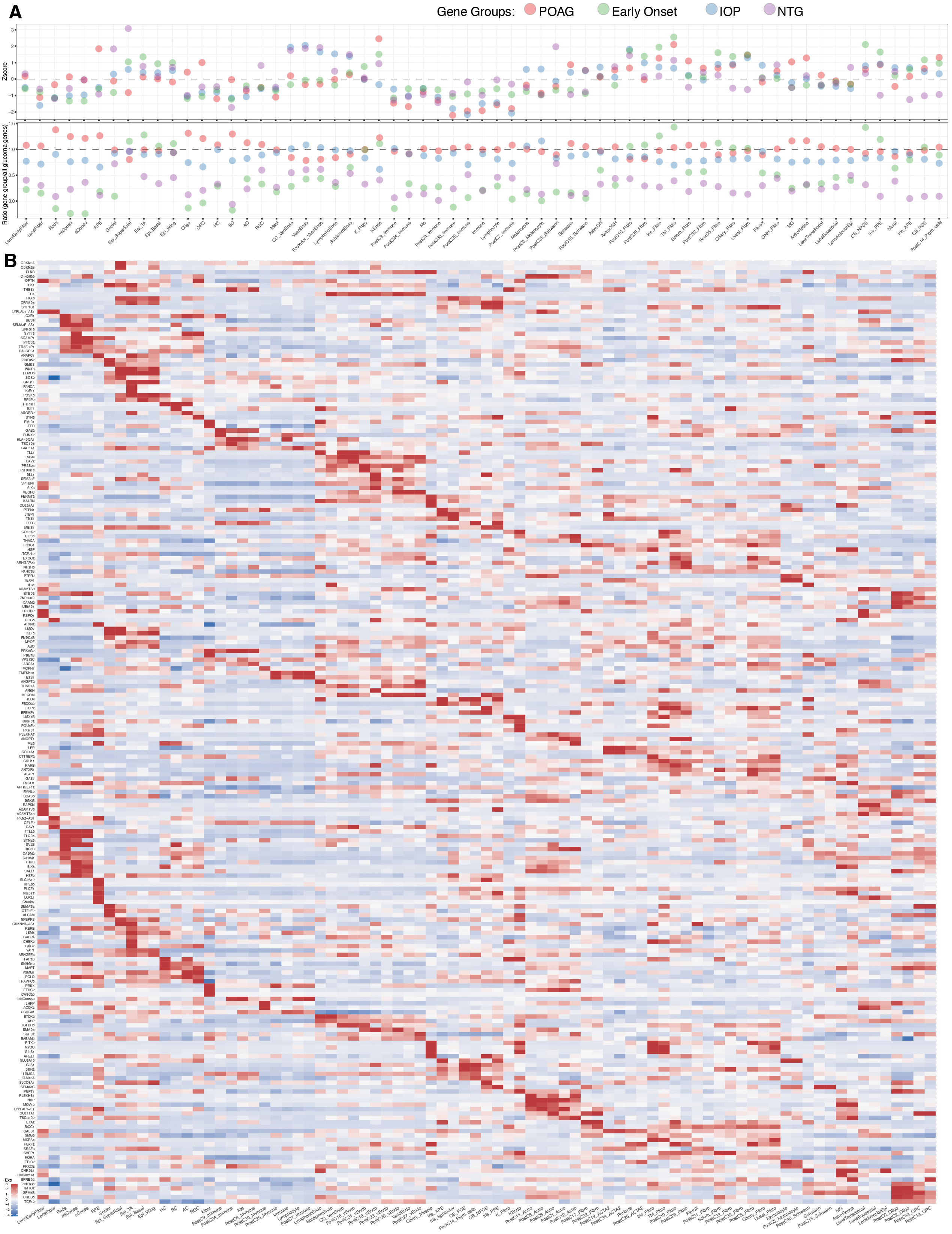
(**A**) Bubble plot showing cell-type specific enrichment z-scores (top) and ratio score (bottom) of groups of genes that have been associated with POAG but not IOP, IOP but not POAG, normal tension glaucoma, or early onset/congenital glaucoma (**B**) Heat map showing expression of all genes associated with primary open angle glaucoma (POAG) and/or intraocular pressure (IOP). Genes are those identified through GWAS analysis (31) (30) and expression is mapped in cell types characterized in the current study and in the recent analysis of the anterior segment van Zyl et al., (9).

## REFERENCES

1. M. Goel, R. G. Picciani, R. K. Lee, S. K. Bhattacharya, Aqueous humor dynamics: a review. The open ophthalmology journal 4, 52 (2010).

2. D. L. Nickla, J. Wallman, The multifunctional choroid. Progress in retinal and eye research 29, 144–168 (2010).

3. J. Forrester, A. Dick, P. McMenamin, F. Roberts, E. Pearlman (2016) Anatomy of the Eye and Orbit, the Eye Basic Sciences in Practice. (Elsevier).

4. J. D. Steinmetz et al., Causes of blindness and vision impairment in 2020 and trends over 30 years, and prevalence of avoidable blindness in relation to VISION 2020: the Right to Sight: an analysis for the Global Burden of Disease Study. The Lancet Global Health 9, e144–e160 (2021).

5. K. B. Freund, D. Sarraf, W. F. Mieler, L. A. Yannuzzi, The retinal Atlas E-book (Elsevier Health Sciences, 2016).

6. D. T. Hartong, E. L. Berson, T. P. Dryja, Retinitis pigmentosa. The Lancet 368, 1795–1809 (2006).

7. H. Quigley, Glaucoma. Lancet. *Glaucoma*. Lancet 377, 1367–1377 (2011).

8. P. Mitchell, G. Liew, B. Gopinath, T. Y. Wong, Age-related macular degeneration. The Lancet 392, 1147–1159 (2018).

9. T. van Zyl et al., Cell atlas of the human ocular anterior segment: Tissue-specific and shared cell types. bioRxiv (2022).

10. T. van Zyl et al., Cell atlas of aqueous humor outflow pathways in eyes of humans and four model species provides insight into glaucoma pathogenesis. Proceedings of the National Academy of Sciences 117, 10339–10349 (2020).

11. Y.-R. Peng et al., Molecular classification and comparative taxonomics of foveal and peripheral cells in primate retina. Cell 176, 1222–1237. e1222 (2019).

12. W. Yan et al., Cell Atlas of The Human Fovea and Peripheral Retina. Scientific Reports 10, 9802(2020).

13. S. W. Lukowski et al., A single-cell transcriptome atlas of the adult human retina. The EMBO journal 38, e100811 (2019).

14. Q. Liang et al., Single-nuclei RNA-seq on human retinal tissue provides improved transcriptome profiling. Nature communications 10, 5743(2019).

15. A. P. Voigt et al., Single-cell transcriptomics of the human retinal pigment epithelium and choroid in health and macular degeneration. Proceedings of the National Academy of Sciences 116, 24100–24107 (2019).

16. M. Menon et al., Single-cell transcriptomic atlas of the human retina identifies cell types associated with age-related macular degeneration. Nature communications 10, 1–9 (2019).

17. L. D. Orozco et al., Integration of eQTL and a single-cell atlas in the human eye identifies causal genes for age-related macular degeneration. Cell reports 30, 1246–1259. e1246 (2020).

18. L. Huang et al., Dynamic human retinal pigment epithelium (RPE) and choroid architecture based on single-cell transcriptomic landscape analysis. Genes & Diseases (2022).

19. J. Collin et al., Single cell RNA sequencing reveals transcriptional changes of human choroidal and retinal pigment epithelium cells during fetal development, in healthy adult and intermediate age-related macular degeneration. Human Molecular Genetics (2023).

20. K. Allison, D. Patel, O. Alabi, Epidemiology of glaucoma: the past, present, and predictions for the future. Cureus 12 (2020).

21. R. G. Strickland, M. A. Garner, A. K. Gross, C. A. Girkin, Remodeling of the Lamina Cribrosa: Mechanisms and Potential Therapeutic Approaches for Glaucoma. International Journal of Molecular Sciences 23, 8068(2022).

22. B. N. Safa, C. A. Wong, J. Ha, C. R. Ethier, Glaucoma and biomechanics. Current Opinion in Ophthalmology 33, 80–90 (2022).

23. J. S. Paula, C. O’Brien, W. D. Stamer, Life under pressure: The role of ocular cribriform cells in preventing glaucoma. Experimental eye research 151, 150–159 (2016).

24. D. J. Calkins, Critical pathogenic events underlying progression of neurodegeneration in glaucoma. Progress in retinal and eye research 31, 702–719 (2012).

25. J. L. Wiggs, L. R. Pasquale, Genetics of glaucoma. Human molecular genetics 26, R21–R27 (2017).

26. N. C. Sears, E. A. Boese, M. A. Miller, J. H. Fingert, Mendelian genes in primary open angle glaucoma. Experimental eye research 186, 107702 (2019).

27. E. R. A. Collantes et al., EFEMP1 rare variants cause familial juvenile-onset open-angle glaucoma. Human mutation 43, 240–252 (2022).

28. J. L. Wiggs, CPAMD8, a new gene for anterior segment dysgenesis and childhood glaucoma. Ophthalmology 127, 767–768 (2020).

29. H. Fu et al., Thrombospondin 1 missense alleles induce extracellular matrix protein aggregation and TM dysfunction in congenital glaucoma. The Journal of Clinical Investigation 132 (2022).

30. P. Gharahkhani et al., Genome-wide meta-analysis identifies 127 open-angle glaucoma loci with consistent effect across ancestries. Nature communications 12, 1–16 (2021).

31. A. P. Khawaja et al., Genome-wide analyses identify 68 new loci associated with intraocular pressure and improve risk prediction for primary open-angle glaucoma. Nature genetics 50, 778–782 (2018).

32. E. G. Hughes, M. E. Stockton, Premyelinating oligodendrocytes: Mechanisms underlying cell survival and integration. Frontiers in Cell and Developmental Biology, 1985(2021).

33. M. Yaqubi et al., Regional and age-related diversity of human mature oligodendrocytes. Glia (2022).

34. S. Jäkel et al., Altered human oligodendrocyte heterogeneity in multiple sclerosis. Nature 566, 543–547 (2019).

35. Z. Liu et al., Specific marker expression and cell state of Schwann cells during culture in vitro. PloS one 10, e0123278 (2015).

36. E. Watanabe, T. Y. Hiyama, R. Kodama, M. Noda, Nax sodium channel is expressed in non-myelinating Schwann cells and alveolar type II cells in mice. Neuroscience letters 330, 109–113 (2002).

37. E. Kimball et al., The role of aquaporin-4 in optic nerve head astrocytes in experimental glaucoma. PloS one 16, e0244123 (2021).

38. L. Muhl et al., Single-cell analysis uncovers fibroblast heterogeneity and criteria for fibroblast and mural cell identification and discrimination. Nature communications 11, 1–18 (2020).

39. C.-C. Deng et al., Single-cell RNA-seq reveals fibroblast heterogeneity and increased mesenchymal fibroblasts in human fibrotic skin diseases. Nature communications 12, 1–16 (2021).

40. F. J. Garcia et al., Single-cell dissection of the human brain vasculature. Nature 603, 893–899 (2022).

41. G. S. Hageman, X. L. Zhu, A. Waheed, W. S. Sly, Localization of carbonic anhydrase IV in a specific capillary bed of the human eye. Proceedings of the National Academy of Sciences 88, 2716–2720 (1991).

42. A. P. Voigt et al., Choroidal endothelial and macrophage gene expression in atrophic and neovascular macular degeneration. Human molecular genetics 31, 2406–2423 (2022).

43. D. Attwell, A. Mishra, C. N. Hall, F. M. O’Farrell, T. Dalkara, What is a pericyte? Journal of Cerebral Blood Flow & Metabolism 36, 451–455 (2016).

44. S.-H. Baek et al., Single Cell Transcriptomic Analysis Reveals Organ Specific Pericyte Markers and Identities. Frontiers in Cardiovascular Medicine 9 (2022).

45. E. A. Winkler et al., A single-cell atlas of the normal and malformed human brain vasculature. Science 375, eabi7377 (2022).

46. T. S. Mitchell, J. Bradley, G. S. Robinson, D. T. Shima, Y.-S. Ng, RGS5 expression is a quantitative measure of pericyte coverage of blood vessels. Angiogenesis 11, 141–151 (2008).

47. L. Ji, S. Xu, H. Luo, F. Zeng, Insights from DOCK2 in cell function and pathophysiology. Frontiers in Molecular Biosciences, 1078(2022).

48. A. Rheinländer, B. Schraven, U. Bommhardt, CD45 in human physiology and clinical medicine. Immunology letters 196, 22–32 (2018).

49. T. Masuda, R. Sankowski, O. Staszewski, M. Prinz, Microglia heterogeneity in the single-cell era. Cell reports 30, 1271–1281 (2020).

50. S. Amor et al., White matter microglia heterogeneity in the CNS. Acta Neuropathologica, 1–17 (2021).

51. A. G. Freud, B. L. Mundy-Bosse, J. Yu, M. A. Caligiuri, The broad spectrum of human natural killer cell diversity. Immunity 47, 820–833 (2017).

52. D. Morgan, V. Tergaonkar, Unraveling B cell trajectories at single cell resolution. Trends in Immunology (2022).

53. R. L. Belote et al., Human melanocyte development and melanoma dedifferentiation at single-cell resolution. Nature Cell Biology 23, 1035–1047 (2021).

54. L. Smith-Thomas et al., Human ocular melanocytes and retinal pigment epithelial cells differ in their melanogenic properties in vivo and in vitro. Current eye research 15, 1079–1091 (1996).

55. Y. Hu et al., Dissecting the transcriptome landscape of the human fetal neural retina and retinal pigment epithelium by single-cell RNA-seq analysis. PLoS Biology 17, e3000365 (2019).

56. Z. Xu et al., A Single-Cell Transcriptome Atlas of the Human Retinal Pigment Epithelium. Frontiers in Cell and Developmental Biology 9 (2021).

57. N. K. Mullin et al., Transcriptomic and chromatin accessibility analysis of the human macular and peripheral retinal pigment epithelium at the single cell level. The American journal of pathology (2023).

58. S. S. Whitmore et al., Transcriptomic analysis across nasal, temporal, and macular regions of human neural retina and RPE/choroid by RNA-Seq. Experimental eye research 129, 93–106 (2014).

59. J. Hahn et al., Evolution of neuronal cell classes and types in the vertebrate retina. bioRxiv (2023).

60. J. E. Craig et al., Multitrait analysis of glaucoma identifies new risk loci and enables polygenic prediction of disease susceptibility and progression. Nature genetics 52, 160–166 (2020).

61. R. L. Radius, M. Gonzales, Anatomy of the lamina cribrosa in human eyes. Archives of ophthalmology 99, 2159–2162 (1981).

62. A. Elkington, C. Inman, P. Steart, R. Weller, The structure of the lamina cribrosa of the human eye: an immunocytochemical and electron microscopical study. Eye 4, 42–57 (1990).

63. H. Lockwood et al., Lamina cribrosa microarchitecture in normal monkey eyes part 1: methods and initial results. Investigative ophthalmology & visual science 56, 1618–1637 (2015).

64. T. Tovar-Vidales, R. J. Wordinger, A. F. Clark, Identification and localization of lamina cribrosa cells in the human optic nerve head. Experimental eye research 147, 94–97 (2016).

65. J. Szeto et al., Regional differences and physiologic behaviors in peripapillary scleral fibroblasts. Investigative ophthalmology & visual science 62, 27–27 (2021).

66. C. Boote et al., Scleral structure and biomechanics. Progress in retinal and eye research 74, 100773 (2020).

67. A. Sridhar et al., Single-cell transcriptomic comparison of human fetal retina, hPSC-derived retinal organoids, and long-term retinal cultures. Cell reports 30, 1644–1659. e1644 (2020).

68. J. Collin et al., A single cell atlas of human cornea that defines its development, limbal progenitor cells and their interactions with the immune cells. The ocular surface 21, 279–298 (2021).

69. P. Gautam et al., Multi-species single-cell transcriptomic analysis of ocular compartment regulons. Nature communications 12, 5675(2021).

70. G. Patel et al., Molecular taxonomy of human ocular outflow tissues defined by single-cell transcriptomics. Proceedings of the National Academy of Sciences 117, 12856–12867 (2020).

71. M. R. Hernandez, F. Igoe, A. H. Neufeld, Cell culture of the human lamina cribrosa. Investigative ophthalmology & visual science 29, 78–89 (1988).

72. N. N. Lopez, A. F. Clark, T. Tovar-Vidales, Isolation and characterization of human optic nerve head astrocytes and lamina cribrosa cells. Experimental eye research 197, 108103 (2020).

73. D. M. Wallace, C. J. O’Brien, The role of lamina cribrosa cells in optic nerve head fibrosis in glaucoma. Experimental eye research 142, 102–109 (2016).

74. R. Rogers, M. Dharsee, S. Ackloo, J. G. Flanagan, Proteomics analyses of activated human optic nerve head lamina cribrosa cells following biomechanical strain. Investigative ophthalmology & visual science 53, 3806–3816 (2012).

75. D. Moore, A. Harris, D. WuDunn, N. Kheradiya, B. Siesky, Dysfunctional regulation of ocular blood flow: A risk factor for glaucoma? Clinical ophthalmology 2, 849–861 (2008).

76. J. Xue et al., Demyelination of the Optic Nerve: An Underlying Factor in Glaucoma? Frontiers in aging neuroscience, 750 (2021).

77. Y. Hao et al., Integrated analysis of multimodal single-cell data. Cell 184, 3573–3587. e3529 (2021).

